# 14-3-3 proteins stabilize actin and vimentin filaments to maintain processes in glomerular podocyte

**DOI:** 10.1101/2023.04.25.538276

**Authors:** Hidenori Yasuda, Yoshiyasu Fukusumi, Ying Zhang, Hiroshi Kawachi

## Abstract

Adaptor protein 14-3-3s have isoform-specific binding partners and roles. We reported 14-3-3β interacts with FKBP12 and synaptopodin to maintain the structure of actin fibers in podocytes. However, differential roles of 14-3-3 isoforms in kidneys are unclear. Herein, we showed that 14-3-3β was dominantly co-localized with FKBP12 in foot processes and was partially co-localized with Par3 at slit diaphragm in podocytes. 14-3-3β interacted with Par3, and FKBP12 bound to 14-3-3β competitively with Par3. Although deletion of 14-3-3β enhanced the interaction of Par3-Par6, it altered actin fiber structure and processes. 14-3-3β and synaptopodin were downregulated in podocyte injury models. 14-3-3σ in podocytes interacted with vimentin in primary processes but not with the actin-associated proteins in foot processes. Deletion of 14-3-3σ altered vimentin fiber structure and processes. 14-3-3σ and vimentin were upregulated in the early phase of podocyte injury models but were decreased in the end stage. Together, the precise localization of 14-3-3β at actin cytoskeleton plays a role in maintaining foot processes and Par complex in podocytes. 14-3-3σ at vimentin cytoskeleton is essential for maintaining primary processes.

## Introduction

The renal glomerular capillary wall (GCW) functions as a barrier, preventing the leakage of plasma protein into the urine. The GCW consists of three layers: an endothelial cell, a glomerular basement membrane (GBM), and a glomerular epithelial cell (podocyte). Podocytes are characterized by a specialized structure of the interdigitating processes covering the outer side of the GBM. The processes of podocyte are composed of two kinds of subcellular compartments, primary processes and secondary processes (foot processes) (1). The cell body projects the primary process, and each primary process protrudes the numerous foot processes. The primary processes composed of intermediate filaments, such as vimentin, and the foot processes composed of actin filaments play an essential role in maintaining the structure of the GCW (2–7). A slit diaphragm connecting the neighboring foot processes is a unique cell-to-cell junction and is reported to be a variant of a tight junction (8). The slit diaphragm functions as a final barrier preventing the leak of plasma proteins into primary urine. Since podocyte dysfunction is a common feature of renal injury in various types of kidney diseases, including minimal-change nephrotic syndrome (MCNS) and focal segmental glomerulosclerosis (FSGS), protecting podocyte from injury is important for slowing or preventing the progression of nephrotic syndrome.

14-3-3 proteins are a ubiquitously expressed family of adaptor proteins initially discovered in brain extracts (9, 10). The 14-3-3 family proteins regulate various signal pathways by binding to many serine or threonine phosphorylated ligands (11–16). In addition, it has been demonstrated that 14-3-3 can bind to non-phosphorylated proteins (11, 17–19). The several mammalian isoforms of 14-3-3 have a variety of expressions in various cell types. Despite exhibiting high sequence homology, it has been identified that 14-3-3 proteins have isoform-specific binding partners and roles (20, 21). For example, 14-3-3σ (also known as stratifin or SFN) is originally characterized as a human mammary epithelial-specific marker that is downregulated in mammary carcinoma cells (22). 14-3-3σ has a unique role in regulating cell cycle progression (23–25) and epithelial differentiation because of their restricted expression, primarily in differentiated epithelial cells (26, 27). We previously identified that 14-3-3β interacts with FK506 binding protein (FKBP) 12 and synaptopodin (Synp) to maintain the structure of actin filaments in podocytes. Tacrolimus (FK506), a widely used immunosuppressive agent, enhances these interactions and ameliorates podocyte injury by restoring FKBP12 at the actin cytoskeleton (28). However, the precise localization and differential role of 14-3-3 isoforms in kidneys are unclear.

The *C. elegans* 14-3-3 homolog, Par5, is required for correct anterior-posterior zygote polarization (29). Other Par proteins, such as Par3 and Par6, have been shown to regulate tight junction formation and epithelial cell polarity. Hurd et al. demonstrated a phosphorylation-dependent mechanism by which 14-3-3 binding regulates the function of the Par3/Par6/aPKC polarity complex in mammalian epithelia (30). Hartleben et al. reported that Par3/Par6/aPKC complex is localized at the slit diaphragm, and the linkage of the major slit diaphragm component nephrin with Par-dependent polarity signaling is essential for establishing the architecture of podocytes (31). We also reported that Par3 interacts with nephrin, and Par6 interacts with another slit diaphragm component ephrinB1, which interacts with nephrin (32, 33). The link of the slit diaphragm complex with the Par complex is essential for maintaining the barrier function of the slit diaphragm. However, whether 14-3-3 proteins are functionally associated with the Par polarity complex in podocytes is unclear.

In the present study, first, we analyzed the distribution of 14-3-3 isoforms in the kidney. 14-3-3β and 14-3-3σ were highly expressed in glomeruli. In glomeruli, 14-3-3β was restricted in podocytes, and 14-3-3σ was expressed in podocytes and mesangial cells. Although 14-3-3β was dominantly co-localized with FKBP12 in the foot processes, a part of 14-3-3β was co-localized with Par3 at the slit diaphragm. Immunoprecipitation (IP) assay showed that 14-3-3β interacted with Par3, and the interaction of 14-3-3β with Par3 was competitive to the interaction with FKBP12. The deletion of 14-3-3β enhanced the interaction of Par3 with Par6 in podocytes. The alteration in filamentous actin (F-actin) structure and deteriorated process formation were observed in the 14-3-3β knockdown (KD) podocytes. 14-3-3σ in podocytes was expressed in the primary processes. 14-3-3σ interacted with vimentin but not with the actin-associated proteins. The alteration in vimentin fiber structure and deteriorated process formation were observed in the 14-3-3σ KD podocytes. 14-3-3β and Synp expression was decreased in the experimental models of both MCNS and FSGS. 14-3-3σ and vimentin expression was clearly increased in the MCNS model but was decreased in the FSGS model. Together, these results suggested that the precise localization of 14-3-3β at the actin cytoskeleton plays a role in maintaining the foot processes and the Par complex in podocytes. In contrast, 14-3-3σ at the vimentin cytoskeleton is essential for maintaining the primary processes.

## Results

### Expression of 14-3-3 isoforms in several organs and tissues

First, mRNA expression of 14-3-3 isoforms was analyzed in mouse cultured podocytes. No difference in mRNA expression of 14-3-3ε, 14-3-3ζ, 14-3-3η, and 14-3-3θ was detected between the undifferentiated and differentiated mouse podocytes. Although mRNA expression of 14-3-3γ in the differentiated mouse podocytes was lower than that in the undifferentiated podocytes, the mRNA expression of 14-3-3σ in the differentiated cells was higher than that in the undifferentiated cells (**Fig EV1A, B**). mRNA and protein expression of 14-3-3β and 14-3-3σ in the differentiated human cultured podocytes was higher than that in the undifferentiated podocytes (**Fig EV1C, D**). It was suggested that 14-3-3β, 14-3-3σ, and 14-3-3γ are related to the differentiation of podocytes. Next, mRNA expression of 14-3-3β, 14-3-3σ, and 14-3-3γ in several major organs was analyzed. mRNA expression of 14-3-3β, 14-3-3σ, and 14-3-3γ was detected in the cerebrum. Expression of 14-3-3β and 14-3-3σ mRNA was detected in several major organs. Expression of 14-3-3γ mRNA in the tissues derived from mesoderm (spleen, kidney, adrenal gland, heart, and skeletal muscle) was higher than in the tissues derived from ectoderm and endoderm. In the skin, 14-3-3σ mRNA was highly expressed compared to other isoforms. In the adrenal gland, 14-3-3β mRNA and 14-3-3γ mRNA were highly expressed compared to 14-3-3σ. In the whole kidney, 14-3-3β mRNA expression was higher than that of 14-3-3σ and 14-3-3γ. 14-3-3β mRNA was equally expressed in all fractions of kidney samples. The expression of 14-3-3σ mRNA in glomeruli was higher than in the cortex and medulla. 14-3-3γ mRNA expression in the cortex was higher than in the glomeruli and medulla (**Fig 1**). These results indicated that 14-3-3γ is highly expressed in renal tubules, and 14-3-3σ is mainly expressed in glomeruli. It was also indicated that 14-3-3β and 14-3-3σ are abundantly expressed in the glomeruli.

**Figure 1.**
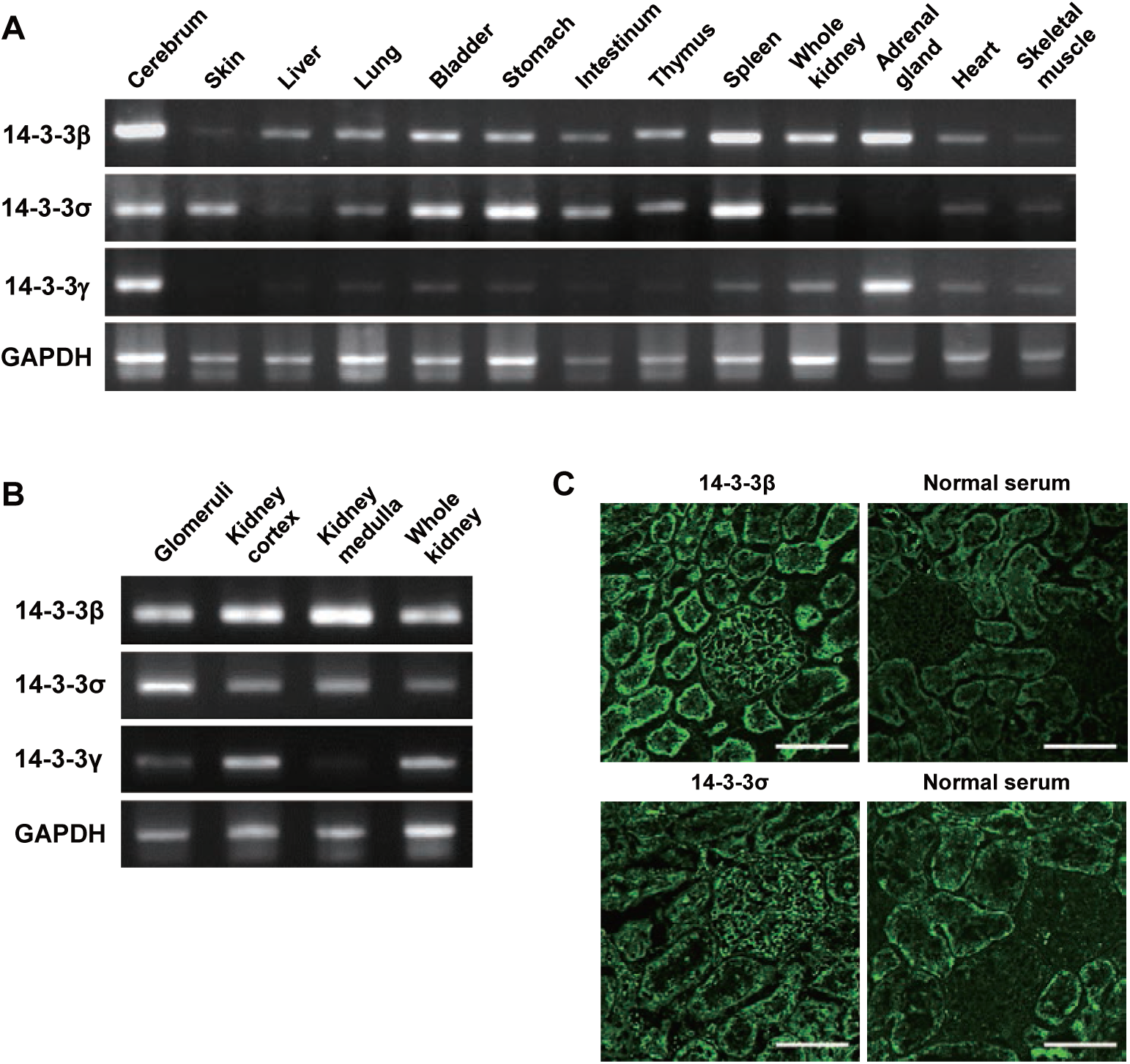
Expression of 14-3-3 isoforms in several organs and tissues. A mRNA expression of 14-3-3β, 14-3-3σ, and 14-3-3γ in the several organs and tissues. mRNA expression of 14-3-3β, 14-3-3σ, and 14-3-3γ was detected in the cerebrum. Expression of 14-3-3β and 14-3-3σ mRNA was detected in several major organs. Expression of 14-3-3γ mRNA in the tissues derived from mesoderm (spleen, kidney, adrenal gland, heart, and skeletal muscle) was higher than that in the tissues derived from ectoderm and endoderm. In the skin, 14-3-3σ mRNA was highly expressed compared to other isoforms. In the adrenal gland, 14-3-3β and 14-3-3γ mRNA was highly expressed compared to 14-3-3σ. B Distribution of 14-3-3β, 14-3-3σ, 14-3-3γ mRNA in the kidney. In the whole kidney, 14-3-3β mRNA expression was higher than that of 14-3-3σ and 14-3-3γ. 14-3-3β mRNA was equally expressed in all fractions of kidney samples. The expression of 14-3-3σ mRNA in glomeruli was higher than that in the cortex and medulla. 14-3-3γ mRNA expression in the cortex was higher than that in the glomeruli and medulla. C Immunofluorescence (IF) findings of 14-3-3β and 14-3-3σ in the rat kidney. The clear positive staining of 14-3-3β and 14-3-3σ was detected in the glomeruli. No positive staining was detected with normal serum control in the glomeruli. Scale bar, 100μm.

### Localization of 14-3-3β and 14-3-3σ in glomeruli and the subcellular localization in podocyte

Next, we analyzed the precise localization of 14-3-3β and 14-3-3σ in glomeruli. A dual-labeling IF study was carried out with glomerular-cell markers: (i) the endothelial cell marker RECA-1; (ii) the mesangial cell marker Thy1.1 and (iii) the podocyte markers (podocalyxin, integrin-α3, nephrin, synaptopodin, and vimentin). 14-3-3β staining was clearly apart from that of RECA-1 and Thy1.1, and no staining co-localized with these markers was detected, indicating that 14-3-3β was restrictedly expressed in the podocytes. Then, to analyze the subcellular localization of 14-3-3β in podocytes, the dual-labeling IF study with podocyte markers was performed. In podocytes, major portions of the 14-3-3β staining were co-stained with Synp, a marker for foot processes, and some portions of the 14-3-3β staining were co-localized with nephrin, a slit diaphragm marker. A part of the 14-3-3β was co-localized with podocalyxin, an apical surface marker, and integrin-α3, a basal surface marker. The staining of 14-3-3β was apart from that of vimentin, a primary process marker (**Fig 2A**). The staining of vimentin was clearly apart from that of Synp (**Fig EV2**). These results indicated that 14-3-3β is dominantly localized in the foot processes and partially localized at the slit diaphragm in podocytes. The staining of 14-3-3σ was clearly apart from that of RECA1, the endothelial cell marker. Some portions of the 14-3-3σ staining were co-localized with Thy.1.1, the mesangial cell marker. In podocytes, major portions of the 14-3-3σ staining were co-stained with vimentin, and some portions of the 14-3-3σ staining were co-stained with podocalyxin and integrin-α3. The 14-3-3σ staining was apart from Synp and nephrin (**Fig 2B**). These results indicated that 14-3-3σ is expressed in mesangial cells and podocytes. 14-3-3σ in podocytes is dominantly localized in primary processes.

**Figure 2.**
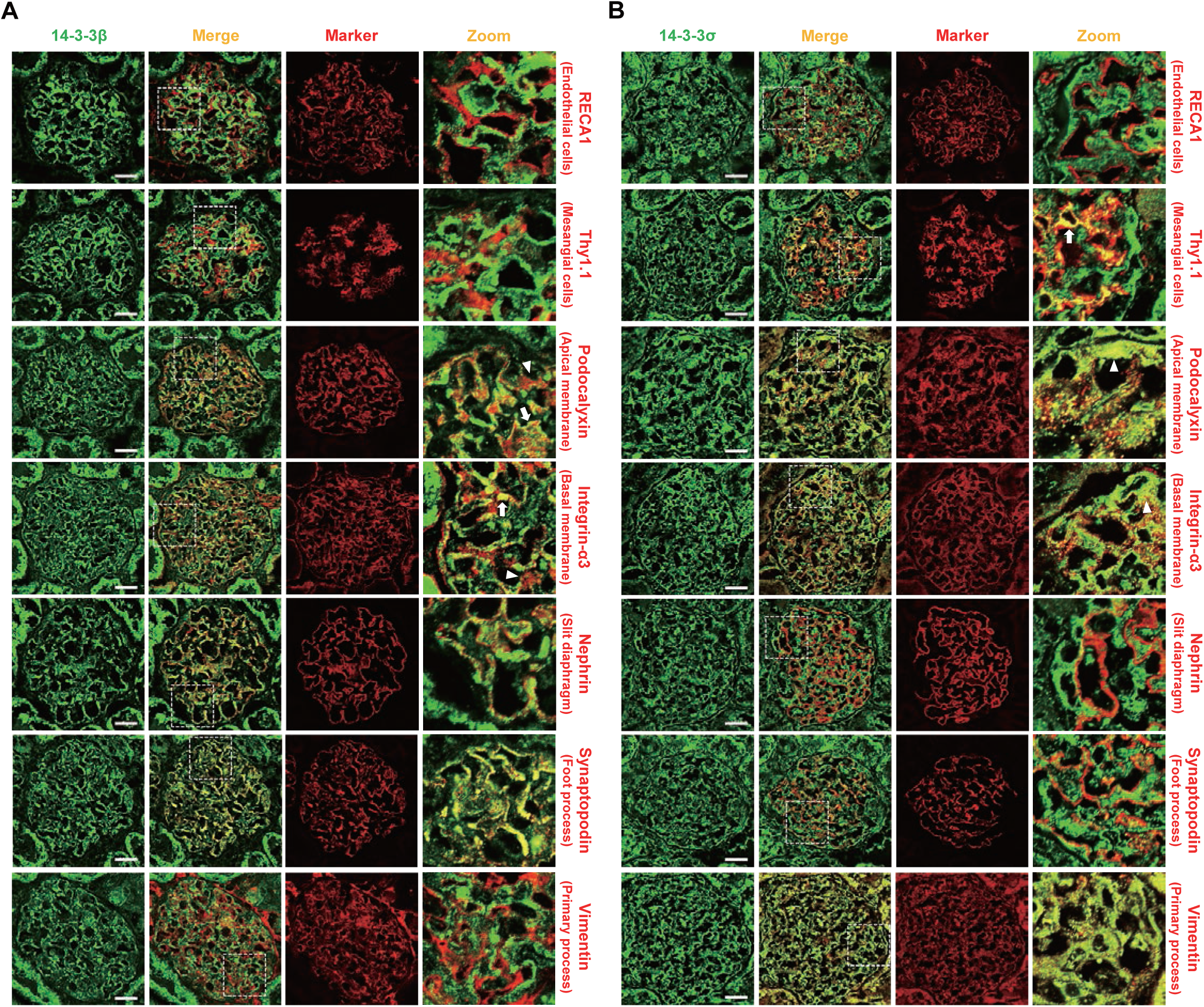
Localization of 14-3-3β and 14-3-3σ in glomeruli and the subcellular localization in podocytes. A Dual-labeling IF findings of 14-3-3β (green) with glomerular cell markers (red) in adult rat glomeruli. The staining of 14-3-3β was clearly apart from that of RECA-1 and Thy1.1, indicating that 14-3-3β is not expressed in endothelial cells or mesangial cells, and 14-3-3β is restricted in podocytes. In podocytes, major portions of the 14-3-3β staining were co-stained with synaptopodin, a foot process marker. Some portions of the 14-3-3β staining were co-localized with nephrin, a slit diaphragm marker. Although a part of the 14-3-3β was co-localized with podocalyxin and integrin-α3 (arrows), apical and basal membrane markers, major portions of the podocalyxin and integrin-α3 staining were apart from the 14-3-3β staining (arrowheads). The 14-3-3β staining was apart from vimentin, a primary process marker. Scale bar, 20μm. B Dual-labeling IF findings of 14-3-3σ (green) with glomerular cell markers (red). The staining of 14-3-3σ was clearly apart from that of RECA-1. A part of 14-3-3σ was co-localized with Thy1.1 (arrow). Major portions of 14-3-3σ were co-stained with vimentin, a primary process marker. Some portions of the 14-3-3σ staining were co-localized with podocalyxin and integrin-α3 (arrowheads). The 14-3-3σ staining was apart from nephrin and synaptopodin. Scale bar, 20μm.

### Interaction of 14-3-3β with Par3 results in the dissociation of Par3 from Par6, and FKBP12 binds to 14-3-3β competitively with Par3 in human cultured podocytes

Next, the association of 14-3-3β with FKBP12 and Par3 was analyzed. In a dual-label IF study with FKBP12 and Par3, 14-3-3β staining was dominantly co-localized with FKBP12. A part of 14-3-3β staining was co-localized with Par3. Some portions of Par3 were apart from 14-3-3β (**Fig 3A**). IP assay with the HEK cells transfected with Par3-myc showed endogenous 14-3-3β interacted with Par3 (**Fig 3B**). The interaction of 14-3-3β with Par3 was also detected in the lysates of cultured podocytes, indicating that 14-3-3β can interact with Par3 in podocytes (**Fig 3C**). The interaction of Par3 and Par6 was detected in the lysates of cultured podocytes (**Fig 3D**). To analyze the effect of 14-3-3β on the Par complex, the interaction of Par3 with Par6 was analyzed in the cultured podocytes treated with 14-3-3β siRNA. Although the expression of Par3 and Par6 was not altered in the podocytes treated with 14-3-3β siRNA, the interaction of Par3 with Par6 was enhanced by the treatment of 14-3-3β siRNA (**Fig 3E-F**). It was indicated that the interaction of 14-3-3β with Par3 resulted in the dissociation of Par3 from Par6 in the cultured podocytes. Furthermore, we analyzed the triadic relationship between 14-3-3β, FKBP12, and Par3. The interaction of 14-3-3β with FKBP12 was detected in the HEK cells transfected with FKBP12-RFP and the cultured podocytes (**Fig 3G**). The amount of endogenous 14-3-3β in the precipitate of Par3 was clearly decreased in the cells transfected with FKBP12, indicating that FKBP12 binds to 14-3-3β competitively with Par3 in podocytes (**Fig 3H**).

**Figure 3.**
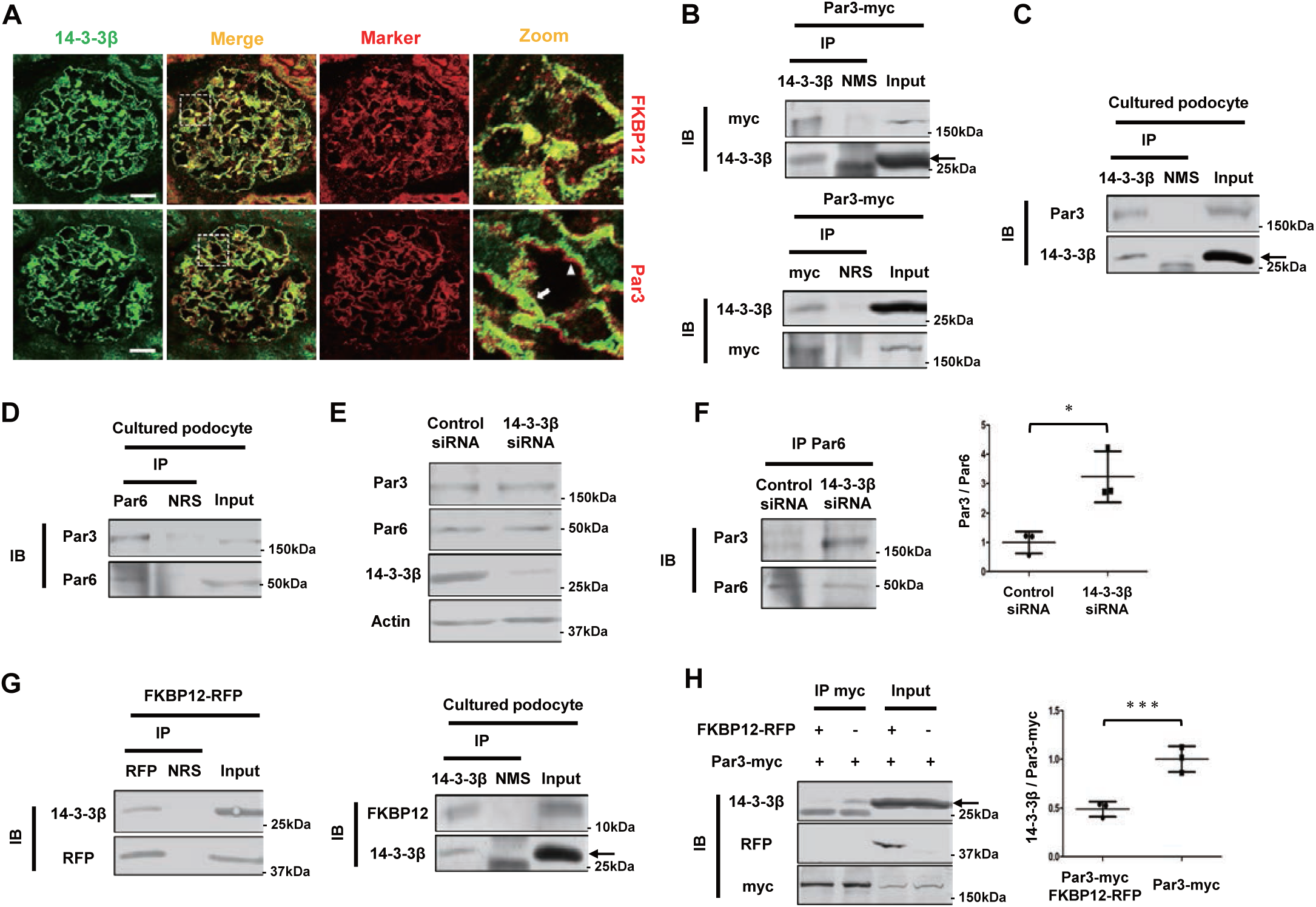
Interaction of 14-3-3β with Par3 results in the dissociation of Par3 from Par6, and FKBP12 binds to 14-3-3β competitively with Par3 in human cultured podocytes. A Dual-labeling IF findings of 14-3-3β (green) with FKBP12 and Par3 (red) in adult rat glomeruli. The major portion of the 14-3-3β staining was co-localized with FKBP12. A part of the 14-3-3β staining was co-stained with Par3 (arrow). Some portions of the Par3 staining were apart from 14-3-3β (arrowhead). Scale bar, 20 μm. B The immunoprecipitation (IP) assay with the HEK293 cells transfected with Par3-myc. The band of Par3 was detected in the precipitate with anti-14-3-3β antibody (upper panel). The endogenous 14-3-3β band was detected in the precipitate with anti-myc antibody (lower panel). 14-3-3β is marked by an arrow. The other 25kDa band is IgG light chain. NMS, normal mouse serum; NRS, normal rabbit serum. C The assay with the human cultured podocytes. Par3 band was detected in the precipitate with anti-14-3-3β antibody. 14-3-3β is marked by an arrow. The other 25kDa band is IgG light chain. NMS, normal mouse serum. D The assay of the interaction of Par3 with Par6 in the cultured podocytes. The band of Par3 was detected in the precipitate with anti-Par6 antibody. NRS, normal rabbit serum. E Western blot finding of Par3, Par6, and 14-3-3β in the cultured podocytes treated with 14-3-3β siRNA. The alteration in protein expression of Par3 and Par6 was not detected in the cells treated with 14-3-3β siRNA. F The assay of the interaction of Par3 with Par6 in the cultured podocyte treated with 14-3-3β siRNA. The band intensity of Par3 was corrected by that of Par6. The date is shown as a ratio relative to the control siRNA-treated cells. The data of three independent experiments are expressed as means ± SD. The amount of Par3 in the Par6 precipitate was clearly increased by the deletion of 14-3-3β (*n* = 3; *p<0.05, *t*-test). G The assay of the interaction of 14-3-3β with FKBP12 in the HEK cells transfected with FKBP12-RFP and the cultured podocytes. 14-3-3β band was detected in the precipitate with anti-RFP antibody (left panel). FKBP12 band was detected in the precipitate with anti-14-3-3β antibody in the cultured podocytes (right panel). 14-3-3β is marked by an arrow. The other 25kDa band is IgG light chain. NMS, normal mouse serum; NRS, normal rabbit serum. H The assay of the interaction of 14-3-3β with Par3 in the cells transfected with FKBP12. The band intensity of 14-3-3β was corrected by that of Par3. The amount of 14-3-3β in the Par3 precipitate was clearly decreased in the cells transfected with FKBP12. The band of FKBP12 was not detected in the precipitate with anti-myc antibody. (means ± SD, *n*=3; ***p<0.001, *t*-test). 14-3-3β is marked by an arrow. The other 25kDa band is IgG light chain.

### 14-3-3σ is a dominant vimentin-binding isoform of 14-3-3 in podocytes

Next, the interaction of 14-3-3σ with vimentin and the actin-associated proteins was analyzed using the HEK cells transfection system. IP assay in the HEK cells co-transfected with 14-3-3σ and vimentin showed that 14-3-3σ interacted with vimentin. The interaction of 14-3-3σ with vimentin was also detected in the lysates of cultured podocytes. The amount of 14-3-3σ in the precipitate of vimentin was clearly higher than that of 14-3-3β, indicating that vimentin predominantly interacts with 14-3-3σ in podocytes (**Fig 4A**). The interaction of 14-3-3σ with FKBP12 was not detected in the lysate of HEK cells co-transfected with 14-3-3σ and FKBP12. The amount of 14-3-3σ in the precipitate of FKBP12 was clearly lower than that of 14-3-3β (**Fig 4B**). The interaction of 14-3-3σ with Synp was almost not detected in the lysate of HEK cells transfected with 14-3-3σ and Synp. The amount of 14-3-3σ in the precipitate of Synp was clearly lower than that of 14-3-3β (**Fig 4C**). These results indicated that 14-3-3β has a high binding affinity with the actin-binding proteins than 14-3-3σ, and 14-3-3σ dominantly interacts with vimentin in podocytes.

**Figure 4.**
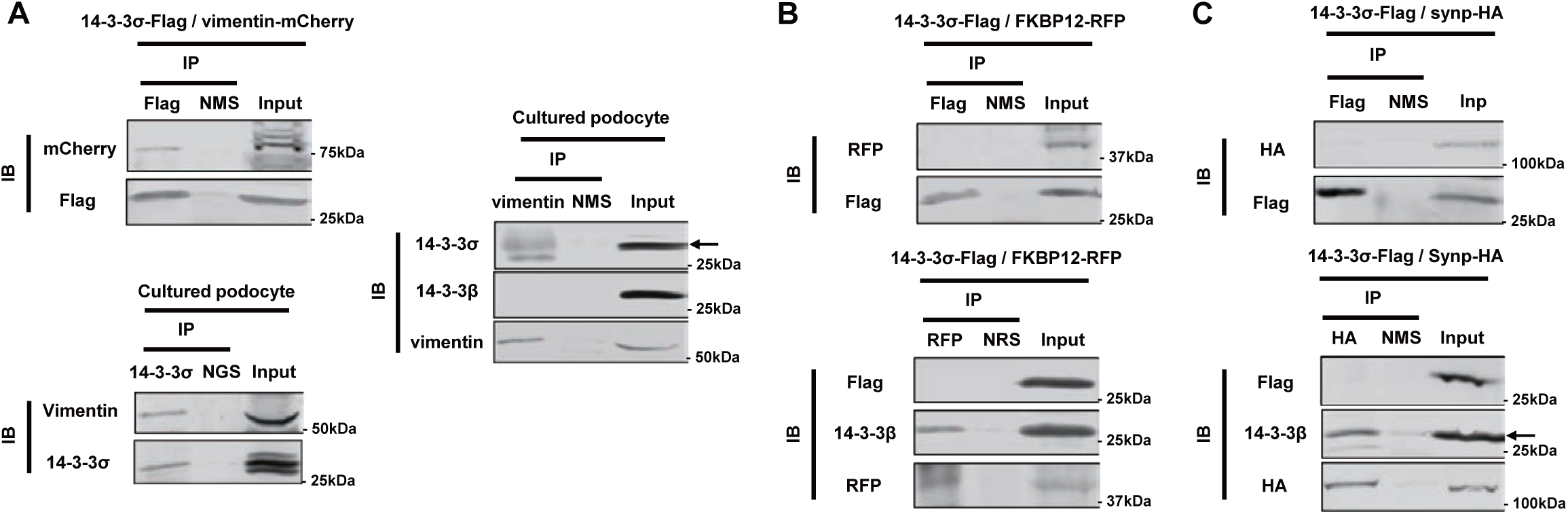
14-3-3σ is a dominant vimentin-binding isoform of 14-3-3 in the podocytes. A The IP assay of the interaction of 14-3-3σ with vimentin in the HEK cells co-transfected with 14-3-3σ-Flag and vimentin-mCherry and in the cultured podocytes. The vimentin band was detected in the precipitate with anti-Flag antibody (left upper panel). The vimentin band was detected in the precipitate with anti-14-3-3σ antibody in the cultured podocytes (left lower panel). The 14-3-3σ band was clearly detected in the precipitate with anti-vimentin antibody in the cultured podocytes. The 14-3-3β band was almost not detected in the precipitate (right panel). 14-3-3σ is marked by an arrow. The other band is nonspecific. NMS, normal mouse serum; NGS, normal goat serum. B The assay of the interaction of 14-3-3σ with FKBP12 in HEK cells transfected with 14-3-3σ-Flag and FKBP12-RFP. The band of FKBP12 was not detected in the precipitate with anti-Flag antibody (upper panel). The 14-3-3σ band was almost not detected in the precipitate with anti-RFP antibody. The 14-3-3β band was detected in the precipitate (lower panel). NMS, normal mouse serum; NRS, normal rabbit serum. C The assay of the interaction of 14-3-3σ with synaptopodin in HEK cells transfected with 14-3-3σ-Flag and synaptopodin-HA (upper panel). The band of synaptopodin was almost not detected in the precipitate with anti-Flag antibody. The 14-3-3σ band was not detected in the precipitate with anti-HA antibody (lower panel). The 14-3-3β band was detected in the precipitate. 14-3-3β is marked by an arrow. The other 25kDa band is IgG light chain. Synp, synaptopodin; NMS, normal mouse serum.

### Deranged F-actin structure, impaired process formation, and increased cell size were detected in 14-3-3β knockdown podocytes

We analyzed the function of 14-3-3β in podocytes using RNA silencing. mRNA expression of 14-3-3β was clearly decreased in the cultured podocytes treated with 14-3-3β siRNA (10.4%). Protein expression of 14-3-3β was decreased in the cells treated with the siRNA (52.7%). mRNA and protein expression of 14-3-3σ was not altered in the cells treated with 14-3-3β siRNA (**Fig 5A**). The large round shape cells were increased, and the proportion of the cells forming processes was decreased by the 14-3-3β siRNA (**Fig 5B-C**). The size of podocytes treated with the siRNA was clearly increased (**Fig 5D**). F-actin staining was clearly decreased in the cells treated with the siRNA, whereas vimentin staining was not altered (**Fig 5C, E**). The proportion of the cells with both stress fibers and vimentin fibers in the cytoplasm was decreased by 14-3-3β siRNA. The proportion of the cells with only vimentin fibers in the cytoplasm was increased in the cells treated with 14-3-3β siRNA (**Fig 5F**). The proportion of the cells with only stress fibers and the cells without stress fibers and vimentin fibers were not altered by the siRNA (**Fig EV3A**). These results indicated that 14-3-3β plays a role in the maintenance of F-actin and the formation of processes in podocytes.

**Figure 5.**
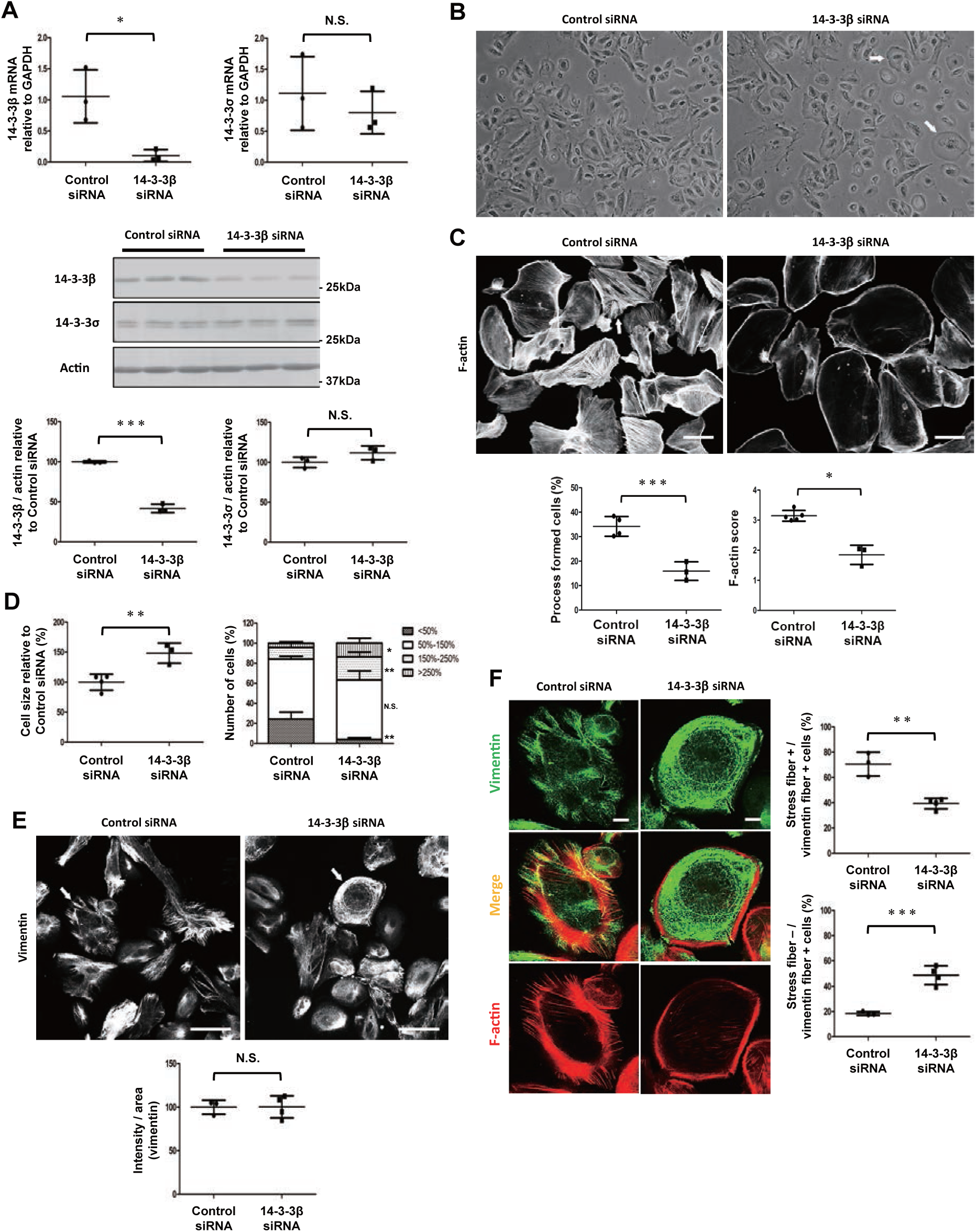
Deranged F-actin structure, impaired process formation, and increased cell size were detected in 14-3-3β knockdown podocytes. A mRNA and protein expression of 14-3-3β and 14-3-3σ in the cultured podocytes treated with 14-3-3β siRNA. GAPDH or actin was analyzed as the internal control. mRNA expression of 14-3-3β was clearly decreased in the podocytes treated with 14-3-3β siRNA. Protein expression of 14-3-3β was decreased in the cells treated with the siRNA. mRNA and protein expression of 14-3-3σ was not altered in the cells treated with siRNA for 14-3-3β. The data are shown as a percentage relative to the control siRNA-treated cells and are expressed as means±SD. (n=3; *p<0.05, ***p<0.005, N.S., non-significant, *t*-test). B Stereomicroscopic findings of the cultured podocytes treated with 14-3-3β siRNA. The round shape cells were increased in the podocytes treated with 14-3-3β (arrows). C F-actin staining in the cultured podocytes treated with 14-3-3β siRNA. The proportion of the cells forming processes (arrow) was decreased in the cells treated with 14-3-3β siRNA. F-actin staining was clearly decreased in the 14-3-3β knockdown podocytes. Scale bar, 100 μm. (n=3-4; ***p<0.001, *t*-test. n=3-5; *p<0.05, *U* test). D Analysis of cell size in the cultured podocytes treated with 14-3-3β siRNA. The size of podocytes treated with 14-3-3β siRNA was clearly increased. The data are shown as a percentage relative to the control siRNA-treated cells and are expressed as means ± SD. The cells <50% in size were decreased in the 14-3-3β knockdown podocytes. The cells 150-250% and >250% in size were increased in the 14-3-3β knockdown podocytes. The size of cells is calculated as a percentage relative to the mean size of the control siRNA-treated cells. (n=3-4; *p<0.05, **p<0.01, *t*-test). E Vimentin staining in the cells treated with 14-3-3β siRNA. The staining intensity of vimentin was not altered in the 14-3-3β knockdown podocytes. Scale bar, 100 μm. (n=3-4; N.S., non-significant, *t*-test). F Dual-labeling IF findings of vimentin and F-actin in the cultured podocyte treated with control siRNA or 14-3-3β siRNA (arrows in E). In the cytoplasm of 14-3-3β knockdown podocytes, stress fibers were clearly decreased, and vimentin fibers were maintained. The proportion of cells with both stress fibers and vimentin fibers in the cytoplasm was decreased in the podocytes treated with 14-3-3β siRNA. The proportion of cells with only vimentin fibers in the cytoplasm was clearly increased in the 14-3-3β knockdown podocytes. Scale bar, 20 μm. (n=3-4; **p<0.01, ***p<0.005, *t*-test).

### Deranged vimentin fiber structure, impaired process formation, and decreased cell size were detected in 14-3-3σ knockdown podocytes

Next, we analyzed the function of 14-3-3σ in podocytes using RNA silencing. mRNA expression of 14-3-3σ was clearly decreased in the cultured podocytes treated with 14-3-3σ siRNA (35.8%). Protein expression of 14-3-3σ was decreased in the cells treated with the siRNA (55.8%). mRNA and protein expression of 14-3-3β was not altered in the cells treated with 14-3-3σ siRNA (**Fig 6A**). The small round shape cells were increased, and the proportion of the cells forming processes was decreased by the 14-3-3σ siRNA (**Fig 6B-C**). The size of podocytes treated with the siRNA was clearly decreased (**Fig 6D**). Although the decrease of F-actin staining was slight, vimentin staining was clearly decreased in the cultured podocytes treated with 14-3-3σ siRNA (**Fig 6C, E**). The proportion of the cells with both stress fibers and vimentin fibers in the cytoplasm was decreased by 14-3-3σ siRNA. The proportion of the cells with only stress fibers in the cytoplasm was increased by 14-3-3σ siRNA (**Fig 6F**). The proportion of the cells with only vimentin fibers was slightly decreased in the cells treated with the siRNA. Alteration in the proportion of the cells without stress fibers and vimentin fibers was not detected (**Fig EV3B**). These findings indicated that 14-3-3σ plays a role in the maintenance of vimentin fibers and the formation of processes in podocytes.

**Figure 6.**
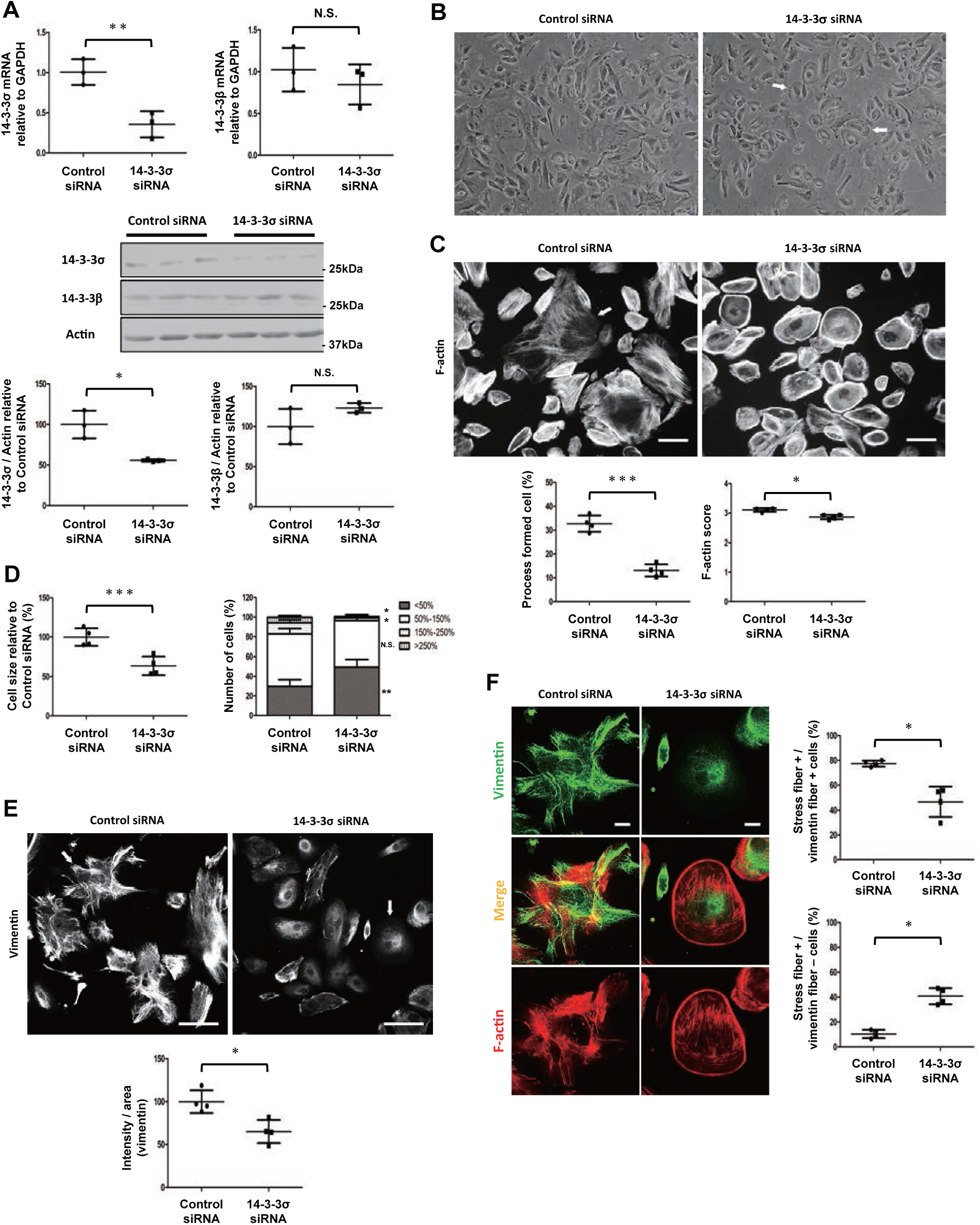
Deranged vimentin fiber structure, impaired process formation, and decreased cell size were detected in 14-3-3σ knockdown podocytes. A mRNA and protein expression of 14-3-3σ and 14-3-3β in the cultured podocytes treated with 14-3-3σ siRNA. GAPDH or actin was analyzed as the internal control. mRNA expression of 14-3-3σ was clearly decreased in the podocytes treated with 14-3-3σ siRNA. Protein expression of 14-3-3σ was decreased in the cells treated with the siRNA. mRNA and protein expression of 14-3-3β was not altered in the cells treated with siRNA for 14-3-3σ. The data are shown as a percentage relative to the control siRNA-treated cells and are expressed as means ± SD. (n=3; *p<0.05, **p<0.01, N.S., non-significant, *t*-test). B Stereomicroscopic findings of the cultured podocytes treated with 14-3-3σ. The round shape cells were increased in the podocytes treated with 14-3-3σ siRNA (arrows). C F-actin staining in the cultured podocytes treated with 14-3-3σ siRNA. The proportion of the cells forming processes (arrow) was clearly decreased in the cells treated with 14-3-3σ siRNA. F-actin staining was slightly decreased in the 14-3-3σ knockdown podocytes. Scale bar, 100 μm. (n=4; ***p<0.001, *t*-test, n=4; *p<0.05, *U* test). D Analysis of cell size in the cultured podocytes treated with 14-3-3σ siRNA. The size of podocytes treated with 14-3-3σ siRNA was clearly decreased. The data are shown as a percentage relative to the control siRNA-treated cells and are expressed as means ± SD. The cells <50% in size were increased in the 14-3-3σ knockdown podocytes. The cells 150-250% and >250% in size were decreased in the 14-3-3σ knockdown podocytes. The size of cells is calculated as a percentage relative to the mean size of the control siRNA-treated cells. (n=4; *p<0.05, **p<0.01, ***p<0.005, *t*-test). E Vimentin staining in the cells treated with 14-3-3σ siRNA. The staining intensity of vimentin was clearly decreased in the 14-3-3σ knockdown podocytes. Scale bar, 100 μm. (n=4; *p<0.05, *t*-test) F Dual-labeling IF findings of vimentin and F-actin in the cultured podocyte treated with control siRNA or 14-3-3σ siRNA (arrows in E). In the cytoplasm of 14-3-3σ knockdown podocytes, vimentin fibers were clearly decreased, and stress fibers were maintained. The proportion of cells with both stress fibers and vimentin fibers in the cytoplasm was decreased in the podocytes treated with 14-3-3σ siRNA. The proportion of cells with only stress fibers in the cytoplasm was clearly increased in the 14-3-3σ knockdown podocytes. Scale bar, 20 μm. (n=4; *p<0.05, *t*-test).

### Expression of 14-3-3β and 14-3-3σ in puromycin-aminonucleoside (PAN)-induced nephropathy and Adriamycin (ADR)-induced nephropathy

Next, we investigated the expression of 14-3-3β and 14-3-3σ in the nephrotic models, PAN nephropathy and ADR nephropathy. PAN nephropathy is accepted to be an experimental model of human MCNS. The staining intensities of nephrin and podocin were decreased on day 10 of PAN nephropathy when proteinuria peaked (**Fig 7A**). mRNA and protein expression of 14-3-3β were decreased on day 10 of PAN nephropathy. The staining intensity of 14-3-3β was also decreased on day 10. In contrast, mRNA and protein expression and staining intensity of 14-3-3σ were clearly increased in the PAN nephropathy (**Fig 7B-D**). ADR nephropathy is accepted to be an experimental model of human FSGS. The staining intensities of nephrin and podocin were clearly decreased on day 28 of ADR nephropathy when severe proteinuria was detected (**Fig 7E**). mRNA and protein expression and staining intensity of 14-3-3β were clearly decreased on day 28 of ADR nephropathy. Although mRNA expression was clearly increased, protein expression and staining intensity of 14-3-3σ were decreased in the ADR nephropathy (**Fig 7F-H**). Dual-label IF study with podocalyxin showed that although the staining of 14-3-3σ in mesangial cells remained, 14-3-3σ staining in podocytes was clearly decreased in the ADR nephropathy (**Fig EV4A**). Furthermore, the expression of 14-3-3β and 14-3-3σ were analyzed in the human cultured podocytes treated with ADR for 14h and 24h. Protein expression of 14-3-3β was decreased in the cultured podocytes treated with ADR for 24h. Protein expression of 14-3-3σ was increased in the podocytes treated with ADR for 14h, whereas the expression of 14-3-3σ was decreased in the podocytes treated with ADR for 24h (**Fig 7I**). The mRNA expression of 14-3-3σ was clearly increased in the cells treated with ADR both for 14h and for 24h (**Fig EV4B**). These results suggested that the downregulation of 14-3-3β is involved in the pathogenesis of nephropathy. It was also suggested that the increase of 14-3-3σ plays a protective role for podocytes in PAN nephropathy, and the downregulation of 14-3-3σ is related to the pathogenesis of ADR nephropathy.

**Figure 7.**
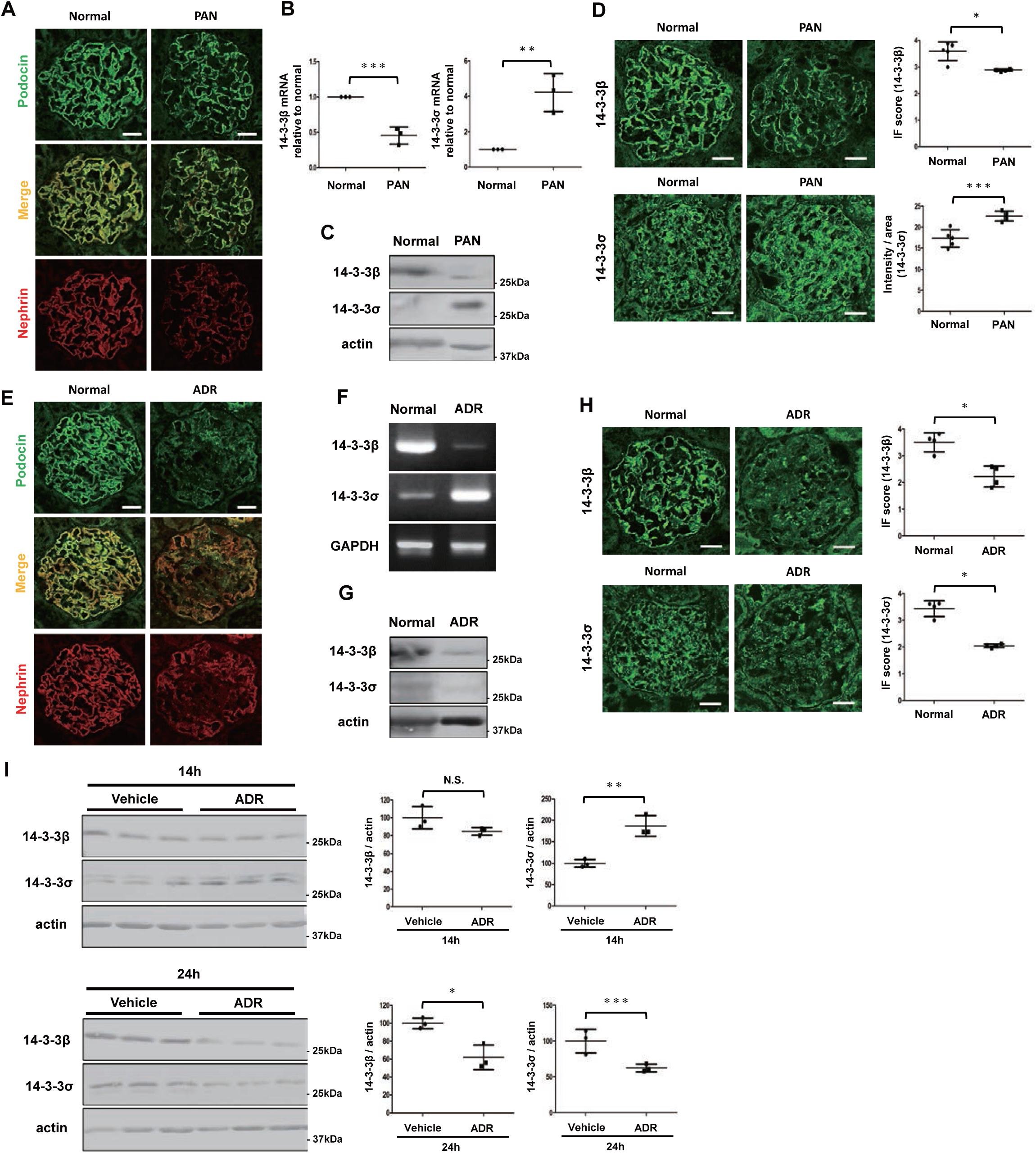
Expression of 14-3-3β and 14-3-3σ in puromycin-aminonucleoside-induced nephropathy and adriamycin-induced nephropathy. A Dual-labeling immunofluorescence (IF) findings of podocin (green) with nephrin (red) on day 10 of puromycin-aminonucleoside (PAN) nephropathy. The staining intensity of nephrin and podocin was clearly decreased on day 10. Scale bar, 20 μm. B mRNA expression of 14-3-3β and 14-3-3σ in PAN nephropathy. The mRNA expression of 14-3-3β was decreased on day 10 of PAN nephropathy. 14-3-3σ mRNA expression was increased on day 10. (means ± SD, *n*=3; **p<0.01, ***p<0.001, *t*-test) C Western blot findings of 14-3-3β and 14-3-3σ on day 10. The protein expression of 14-3-3β was decreased on day 10. The protein expression of 14-3-3σ was clearly increased. D IF findings of 14-3-3β and 14-3-3σ on day 10 of PAN nephropathy. The staining intensity of 14-3-3β was decreased. The staining score is shown as means ± SD (*n*=5; *p<0.05, *U* test). The staining intensity of 14-3-3σ was increased. The staining intensity par area is shown as means ± SD. Scale bar, 20 μm. (*n*=5; ***p<0.001, *t*-test). E Dual-IF findings of podocin (green) and nephrin (red) on day 28 in adriamycin (ADR) nephropathy. The staining intensity of nephrin and podocin was clearly decreased on day 28 of ADR nephropathy. Scale bar, 20 μm. F mRNA expression of 14-3-3β and 14-3-3σ in ADR nephropathy. mRNA expression of 14-3-3β was decreased on day 28 of ADR nephropathy. 14-3-3σ mRNA expression was increased on day 28. G Western blot findings of 14-3-3β and 14-3-3σ on day 28. The protein expression of 14-3-3β and 14-3-3σ was decreased on day 28. H IF findings of 14-3-3β and 14-3-3σ on day 28 of ADR nephropathy. The staining intensity of 14-3-3β and 14-3-3σ was decreased. Scale bar, 20 μm. (*n*=4; * p <0.05, *U* test). I Western blot finding of 14-3-3β and 14-3-3σ in the human cultured podocytes treated with ADR for 14 h and 24 h. The protein expression of 14-3-3β was decreased in the cultured podocytes treated with 24 h. The protein expression of 14-3-3σ was increased in the cultured podocyte treated with 14h and was decreased in the cells treated with 24 h. (*n*=3; *p<0.05, **p<0.01, ***p<0.001, N.S., non-significant, *t*-test).

### Deranged F-actin and vimentin fiber structure and impaired process formation were detected in 14-3-3β and 14-3-3σ double knockdown podocytes

Next, we analyzed the effect of double gene silencing for 14-3-3β and 14-3-3σ on the podocytes. mRNA and protein expression were decreased in the cultured podocytes treated with both 14-3-3β siRNA and 14-3-3σ siRNA (**Fig 8A**). The round shape cells were increased, and the proportion of the cells forming processes was decreased in the podocytes treated with 14-3-3β and 14-3-3σ siRNA (**Fig 8B-C**). The size of podocytes treated with these siRNAs was not altered (**Fig 8D**). F-actin and vimentin staining was clearly decreased by the 14-3-3β and 14-3-3σ siRNA (**Fig 8C, E**). The proportion of the cells with both stress fibers and vimentin fibers in the cytoplasm was decreased by the 14-3-3β and 14-3-3σ siRNA. The proportion of the cells with only stress fibers or only vimentin fibers was increased in the cells treated with these siRNAs. The proportion of the cells without both stress fibers and vimentin fibers was clearly increased by these siRNAs (**Fig 8F**). These findings suggested that the downregulation of 14-3-3β and 14-3-3σ in ADR-induced podocyte injury disrupts the structure of actin and vimentin fibers.

**Figure 8.**
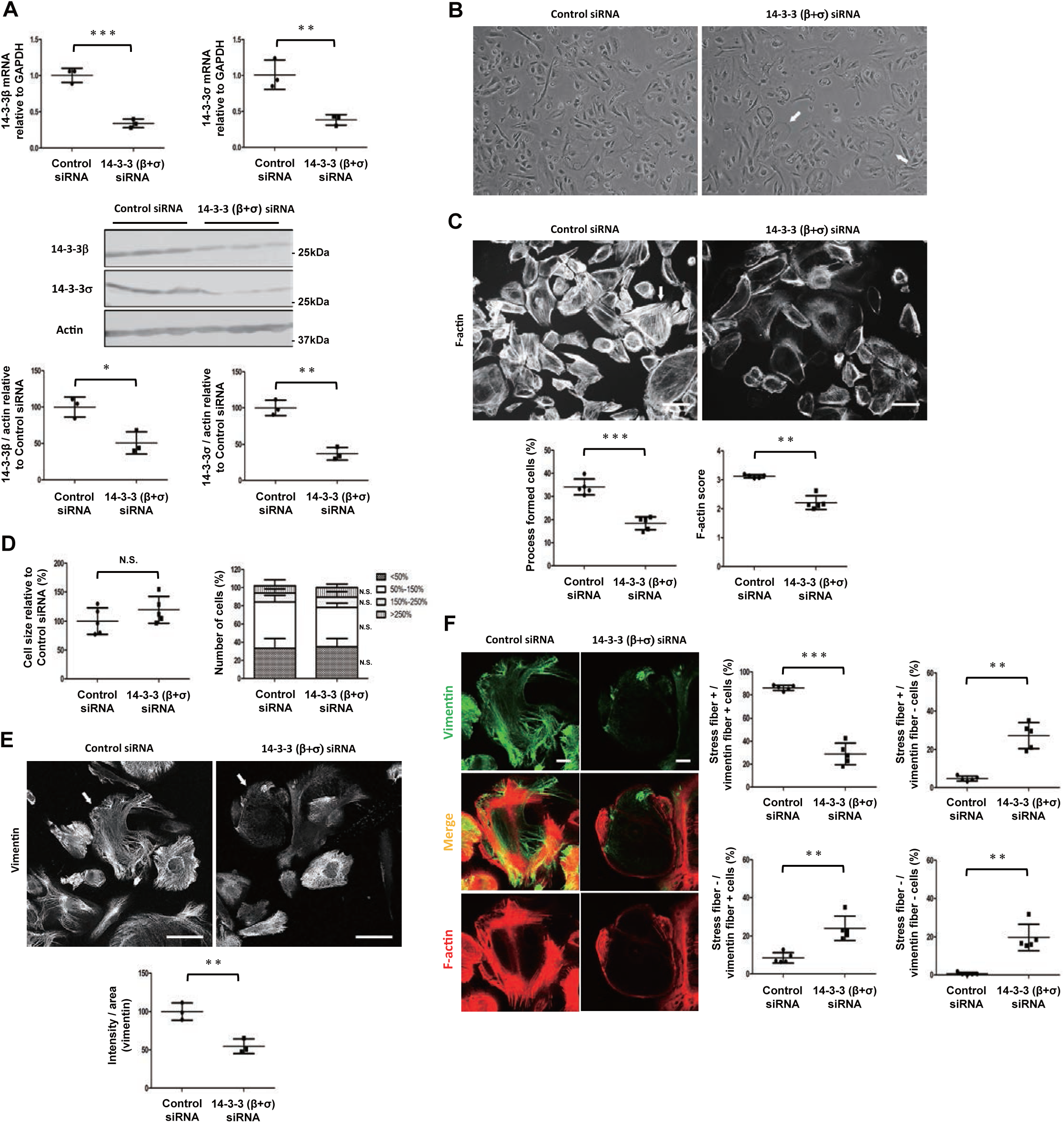
Deranged F-actin and vimentin fiber structure and impaired process formation were detected in 14-3-3β and 14-3-3σ double knockdown podocytes. A mRNA and protein expression of 14-3-3β and 14-3-3σ in the cultured podocytes treated with 14-3-3β and 14-3-3σ siRNA. (*n*=3; *p<0.05, **p<0.01, ***p<0.001, *t*-test). B Stereomicroscopic findings of the cultured podocytes treated with 14-3-3β and 14-3-3σ siRNA. The round shape cells were increased in the podocytes treated with 14-3-3β and 14-3-3σ siRNA (arrows). C F-actin staining in the cultured podocytes treated with 14-3-3β and 14-3-3σ siRNA. The proportion of the cells forming processes (arrow) was decreased in the cells treated with 14-3-3β and 14-3-3σ siRNA. F-actin staining was clearly decreased in the 14-3-3σ knockdown podocytes. Scale bar, 100 μm. (n=5; ***p<0.001, *t*-test, n=5; **p<0.05, *U* test). D Analysis of cell size in the cultured podocytes treated with 14-3-3β and 14-3-3σ siRNA. The alteration in size of podocytes treated with 14-3-3β and 14-3-3σ siRNA was not detected. The data are shown as a percentage relative to the control siRNA-treated cells and are expressed as means ± SD. The distribution of the cell size in the 14-3-3β and 14-3-3σ knockdown podocytes was not altered. (n=5; N.S., non-significant, *t*-test). E Vimentin staining in the cells treated with 14-3-3β and 14-3-3σ siRNA. The staining intensity of vimentin was clearly decreased in the 14-3-3β and 14-3-3σ knockdown podocytes. Scale bar, 100 μm. (n=3; **p<0.01, *t*-test). F Dual-labeling IF findings of vimentin and F-actin in the cultured podocyte treated with control siRNA or 14-3-3β and 14-3-3σ siRNA (arrows in E). Stress fibers and vimentin fibers were clearly decreased in the cytoplasm of 14-3-3β and 14-3-3σ knockdown podocytes. The proportion of cells with both stress fibers and vimentin fibers in the cytoplasm was decreased in the podocytes treated with 14-3-3β and 14-3-3σ siRNA. The proportion of cells with only stress fibers or only vimentin fibers in the cytoplasm was clearly increased in the 14-3-3β and 14-3-3σ knockdown podocytes. The proportion of cells without both stress fibers and vimentin fibers in the cytoplasm was increased in the double knockdown podocytes. Scale bar, 20 μm. (n=5; **p<0.01, ***p<0.005 *t*-test).

### Primary processes and foot processes were deteriorated in ADR-induced nephropathy

Next, the expression of the primary process and foot process markers was analyzed in podocyte injury models. The round shape cells clearly increased in the cultured podocytes treated with ADR for 24h (**Fig 9A**). Although the alteration in the proportion of cells forming processes was not detected in the podocytes treated with ADR for 14h, that was clearly decreased in the cells treated with ADR for 24h. F-actin staining was decreased in the podocytes treated with ADR for 14h and 24h (**Fig 9B**). Vimentin staining was increased in the podocytes treated with ADR for 14h but clearly decreased in the cells treated with ADR for 24h (**Fig 9C**). Although stress fibers in the cytoplasm were decreased, vimentin fibers were increased in the podocytes of ADR 14h. In the cells treated with ADR for 24h, both stress fibers and vimentin fibers in the cytoplasm were clearly decreased (**Fig 9D**). These results suggested that although the foot processes are deranged, the primary processes are maintained or upregulated in the early phase of ADR-induced podocyte injury, subsequently, both the foot processes and the primary processes are deteriorated in the end phase. Furthermore, we analyzed the expression of vimentin and Synp in PAN nephropathy and ADR nephropathy. Although Synp staining was decreased, vimentin staining was clearly increased on day 10 of the PAN nephropathy (**Fig 9E**). The staining of Synp and vimentin was decreased on day 28 of the ADR nephropathy (**Fig 9F**). These findings indicated that although the foot processes are deranged, the primary processes are upregulated in the PAN nephropathy. It was also indicated that both foot processes and primary processes deteriorated in ADR nephropathy.

**Figure 9.**
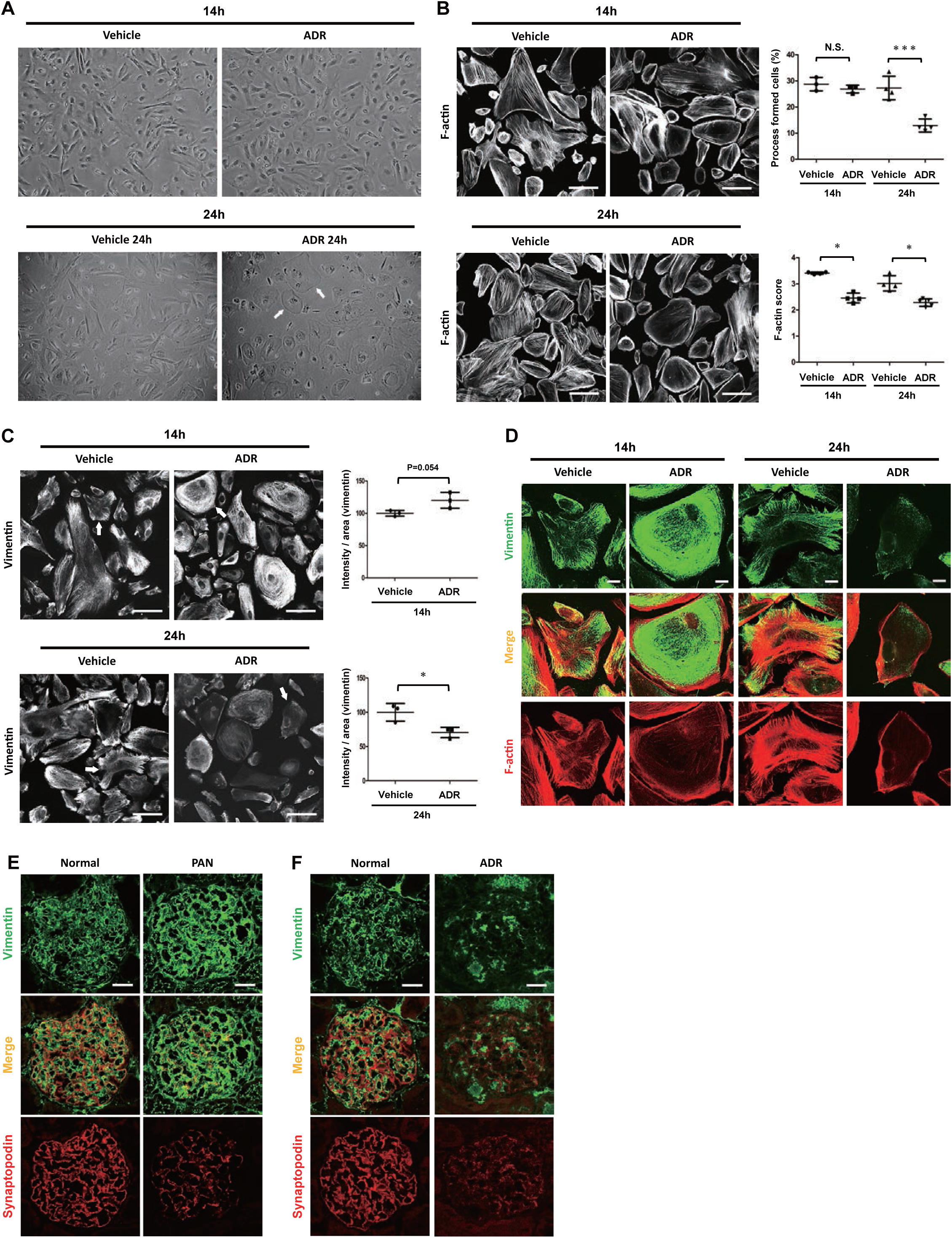
Primary processes and foot processes were deteriorated in ADR-induced nephropathy. A Stereomicroscopic findings of the cultured podocytes treated with ADR for 14h and 24h. The round shape cells were clearly increased in the podocytes treated with ADR for 24h (arrows). B F-actin staining in the cultured podocytes treated with ADR 14h and 24h. The proportion of the cells forming processes was decreased in the cells treated with ADR for 24h. The alteration in the proportion of the cells forming processes was not detected in the cells treated with ADR for 14h. F-actin staining was decreased in the podocytes treated with ADR for 14h and 24h. Scale bar, 100 μm. (n=3-4; ***p<0.001, N.S., non-significant, *t*-test, n=4; *p<0.05, *U* test). C Vimentin staining in the cultured podocytes treated with ADR for 14h and 24h. The staining intensity of vimentin was increased in the cells treated with ADR 14h. The staining intensity of vimentin was clearly decreased in the cells treated with ADR 24h. Scale bar, 100 μm. (n=3; *p<0.05, *t*-test) D Dual-labeling IF staining of vimentin with F-actin in the cultured podocyte treated with ADR 14h and 24h (arrows in C). The decrease of stress fibers and the increased vimentin fibers in the cytoplasm were observed in the cells treated with ADR 14h. Both vimentin and F-actin staining was clearly decreased in the cells treated with ADR 24h. Scale bar, 20 μm. E Dual-labeling IF staining of vimentin with synaptopodin on day 10 of PAN-induced nephropathy. The staining intensity of vimentin was clearly increased in PAN nephropathy. Synaptopodin staining was clearly decreased in the PAN nephropathy. Scale bar, 20 μm. F Dual-labeling IF of vimentin with synaptopodin on day 28 of ADR nephropathy. The staining intensity of vimentin and synaptopodin was decreased in ADR nephropathy. Scale bar, 20 μm.

## Discussion

It has been reported that 14-3-3 isoforms have isoform-specific binding partners and roles (20). We previously identified that 14-3-3β interacts with the actin-associated proteins, FKBP12 and Synp, to maintain the structure of actin fibers in podocytes (28). It has also been reported that 14-3-3 regulates the function of Par3/Par6/aPKC complex in mammalian epithelia (30). The precise localization and differential roles of 14-3-3 isoforms in kidneys are unclear. Whether 14-3-3 proteins are functionally associated with the Par polarity complex in podocytes is also unclear. In the present study, we investigated the expression and function of 14-3-3 isoforms in kidneys and analyzed the differential roles in the pathogenesis of nephrotic syndromes.

First, we analyzed the expression of 14-3-3 isoforms in several major organs. In the kidney, 14-3-3γ was highly expressed in the cortex than that in the glomeruli and the medulla, indicating 14-3-3γ is dominantly expressed in renal tubules. 14-3-3β was equally expressed in glomeruli, cortex, and medulla, indicating 14-3-3β is broadly expressed in glomeruli and tubules. Although the expression of 14-3-3σ in the kidney was lower than that in other organs, 14-3-3σ was highly expressed in glomeruli. IF study showed 14-3-3β and 14-3-3σ were abundantly expressed in glomeruli (**Fig 1**). Dual-labeling IF studies with glomerular cell markers revealed that 14-3-3β is exclusively expressed in podocytes, whereas 14-3-3σ is expressed in podocytes and a part of mesangial cells (**Fig 2**). The difference in the distribution of 14-3-3β, 14-3-3σ, and 14-3-3γ suggests cell-specific roles of these 14-3-3 isoforms in the kidney. These results suggested that 14-3-3β and 14-3-3σ function in podocytes.

Next, we analyzed the subcellular localization of 14-3-3β and 14-3-3σ in podocytes. The dual-labeling IF study with podocyte markers showed that 14-3-3β is dominantly co-localized with Synp, the actin-associated protein of foot processes. A part of 14-3-3β was co-localized with nephrin, a slit diaphragm component. In contrast, 14-3-3σ is dominantly co-localized with vimentin, the intermediate filament of primary processes (**Fig 2**). 14-3-3β and 14-3-3σ were highly expressed in the differentiated human cultured podocytes with processes (**Fig EV1**). In the present study, we identified the precise localization of 14-3-3β and 14-3-3σ in glomeruli. These results suggest that 14-3-3β interacts with not only the actin-associated proteins but also the slit diaphragm components. It is also suggested that 14-3-3σ interacts with vimentin and is involved in maintaining the primary processes.

Next, we analyzed the interaction of 14-3-3β with actin-associated protein FKBP12 and Par polarity protein Par3. We previously identified the interaction of 14-3-3β with FKBP12 (28). Dual-label IF study revealed that 14-3-3β is dominantly co-localized with FKBP12. Although a part of 14-3-3β is co-localized with Par3, a Par polarity protein that interacts with nephrin, some portions of Par3 are apart form 14-3-3β in the normal glomeruli. The IP assay with the HEK transfection system revealed that 14-3-3β interacts with Par3, and the interaction was confirmed by the analysis with cultured podocytes. Par3 interacted with Par6 in the cultured podocytes. The link of the Par3-Par6 complex with the slit diaphragm complex is essential for maintaining the barrier function of the slit diaphragm (32). Benton et al. reported that 14-3-3 binding to Bazooka (*Drosophila* Par3 homolog) inhibits the formation of Bazooka/Par6/aPKC polarity complex in *Drosophila* (34). Hurd et al. reported that overexpression of 14-3-3 results in a severe disruption of polarity in mammalian epithelia (30). In the present study, although the expression of Par3 and Par6 was not altered, the interaction of Par3 with Par6 was clearly enhanced in the 14-3-3β-deleted cultured podocytes. This result indicates that the interaction of 14-3-3β with Par3 disrupts the Par complex at the slit diaphragm in podocytes. Furthermore, the IP assay with the HEK transfection system revealed that the interaction of 14-3-3β with Par3 is competitive to the interaction with FKBP12 (**Fig 3**). These results suggest that the binding of 14-3-3β with the actin-associated protein FKBP12 is essential for the maintenance of Par complex at the slit diaphragm. We have reported that tacrolimus, a widely used immunosuppressive agent, enhances the interaction of FKBP12 with 14-3-3β and Synp (28). It is plausible that the effect of tacrolimus on the competitive interaction of 14-3-3β with Par3 also participates in the podocyte protective effect.

Next, we analyzed the interaction of 14-3-3σ with vimentin and the actin-associated proteins. The IP assay with the HEK transfection system revealed that 14-3-3σ interacts with vimentin, and the interaction of 14-3-3σ with vimentin was confirmed by the assay with cultured podocytes (**Fig 4**). Tzivion et al. reported that the vimentin amino-terminal head domain (amino acids 1-96) is necessary for binding to 14-3-3, and the stable 14-3-3 dimerization is required for the binding to two sites on a single vimentin polypeptide (35). Wang et al. also reported the interaction of 14-3-3 with vimentin filament (36). We previously reported that FKBP12 interacts with 14-3-3β and Synp and plays a role in the maintenance of the expression of 14-3-3β (28). Faul et al. also reported that Synp is stabilized through binding with 14-3-3β in podocytes, and the 14-3-3β-Synp binding maintains stress fibers in podocytes (37, 38). In the present study, IP assay with the cultured podocytes revealed that the interaction of 14-3-3σ with vimentin is clearly higher than the interaction of 14-3-3β. Furthermore, the HEK transfection system revealed that the binding affinity of 14-3-3β with the actin-associated proteins, FKBP12 and Synp, is clearly higher than that of 14-3-3σ (**Fig 4**). These results indicated that the difference in binding affinity to the actin-associated proteins or vimentin is involved in the differential localization of 14-3-3β and 14-3-3σ in podocytes. It was suggested that 14-3-3β and 14-3-3σ play differential roles in maintaining cytoskeletal filaments in podocytes. To investigate this hypothesis, 14-3-3β or 14-3-3σ KD podocytes were analyzed. Alteration of F-actin, deteriorated process formation, and increased cell size were observed in the 14-3-3β KD podocytes (**Fig 5**). In contrast, alteration of vimentin fibers, deteriorated process formation, and decreased cell size were observed in the 14-3-3σ KD cells (**Fig 6**). These findings indicated that 14-3-3β and 14-3-3σ are essential for maintaining the processes in podocytes. 14-3-3β and 14-3-3σ play roles in maintaining F-actin and vimentin fibers, respectively. This is the first report of the differential function of 14-3-3β and 14-3-3σ for maintaining cytoskeletal filaments in podocytes.

To elucidate the pathogenic significance of 14-3-3β and 14-3-3σ in nephrotic models, we investigated the expression of 14-3-3β and 14-3-3σ in PAN-induced or ADR-induced podocyte injury. The expression of 14-3-3β was decreased, and 14-3-3σ expression was clearly increased in PAN nephropathy, a mimic of MCNS. The expression of both 14-3-3β and 14-3-3σ was decreased in ADR nephropathy, a mimic of FSGS (**Fig 7**). Mass et al. proposed that MCNS is the early stage of FSGS (39). FSGS is one of the most common primary glomerulonephropathies (40). FSGS patients are cortisone-resistant, with a tendency to progress to end-stage renal disease (41). In the present study, although 14-3-3σ mRNA was constantly increased, protein expression of 14-3-3σ was increased in the early phase and then decreased in the end phase of cultured podocytes treated with ADR (**Fig 7**). This result suggests that the alteration of 14-3-3σ in the early phase of ADR-treated cultured podocytes represents that in the PAN nephropathy. It was also suggested that the downregulation of both 14-3-3β and 14-3-3σ participates in the pathogenesis of ADR nephropathy. To investigate this hypothesis, the structure of F-actin and vimentin fibers in 14-3-3β and 14-3-3σ double KD podocytes and ADR-treated podocytes were analyzed. Expectedly, alteration of F-actin and vimentin fibers, deteriorated process formation, and severe morphological alteration were observed in the 14-3-3β and 14-3-3σ double KD cells (**Fig 8**). In the cultured podocytes treated with ADR for 14h, although the F-actin structure was altered, vimentin fibers in the cytoplasm were upregulated, and process formation was maintained. In the cultured podocytes with ADR for 24h, alteration of both F-actin and vimentin fibers and deteriorated process formation were observed. These results suggested that the downregulation of 14-3-3β and 14-3-3σ participates in the deterioration of primary and foot processes in podocyte injury. It is plausible that the upregulation of 14-3-3σ is a protective response to podocyte injury. Furthermore, the expression of vimentin, as a primary process marker, and Synp, as a foot process marker, was analyzed in PAN and ADR nephropathy. Vimentin is a widely distributed intermediate filament protein in mesenchyme-derived cells and is also used as a marker of the epithelial-to-mesenchymal transitions (42, 43). Podocytes protrude a rich elastic network of vimentin fibers in the primary processes that are important in providing structural stability (44, 45). The actin-associated protein Synp is highly expressed in the foot processes of podocytes (46). In podocytes, gene silencing of Synp causes the loss of F-actin and RhoA activity, which indicates that Synp is one of the critical molecules in podocyte (37, 47, 48). Although Synp expression was decreased, vimentin expression was clearly increased in PAN nephropathy. In contrast, the expression of both Synp and vimentin was decreased in ADR nephropathy (**Fig 9**). Zou et al. also reported that the expression of vimentin and nestin, intermediate filaments localized at the primary processes, are increased in PAN nephropathy (5). Su et al. reported that nestin expression was reduced in FSGS patients (49). Dijkman et al. reported that vimentin is expressed in the periglomerular area in glomeruopathic sclerosis of MCNS patients (50). Ichii et al. also reported that Synp and vimentin were decreased in the glomeruli of autoimmune glomerulonephritis model mice with progressive disease (51). All of these findings suggest that 14-3-3β and 14-3-3σ maintain the primary and foot processes by stabilizing actin and vimentin cytoskeleton in podocytes, and the downregulation of these 14-3-3 proteins participates in the pathogenesis of FSGS. Recently, Wang et al. also reported that the expression of 14-3-3σ in kidneys is upregulated in the acute kidney injury (AKI) model mice caused by ischemia-reperfusion (I/R) (52). However, they reported the upregulation of 14-3-3σ in renal tubules activated necroptosis in cisplatin-or I/R-induced AKI models, suggesting that 14-3-3σ may play distinct roles in the tubular epithelial cells without vimentin.

In conclusion, 14-3-3β in glomeruli is restrictedly expressed in the podocytes. 14-3-3β in podocytes is expressed at the foot processes and a part of the slit diaphragm. 14-3-3β interacts with FKBP12 and Synp in the foot processes. Although 14-3-3β can interact with Par3, the interaction of 14-3-3β with Par3 is competitive to the interaction with the actin-associated protein FKBP12. The interaction of 14-3-3β with Par3 disrupts the Par3-Par6 complex in podocytes. In the pathological state, 14-3-3β is downregulated, and the decrease of 14-3-3β alters the structure of actin fibers. In contrast, 14-3-3σ in glomeruli is expressed in the podocytes and the mesangial cells. 14-3-3σ in podocytes is expressed at the primary processes. The binding affinity of 14-3-3σ with the actin-associated proteins is lower than that of 14-3-3β. 14-3-3σ predominantly interacts with vimentin. In the pathological state, 14-3-3σ is transiently upregulated in the early stage, then downregulated in the end stage. The decrease of 14-3-3σ altered the structure of vimentin fibers. The expression of both 14-3-3β and 14-3-3σ is downregulated, and both primary and foot processes are deteriorated in the end stage of podocyte injury, such as FSGS. In the present study, we demonstrated that the precise localization of 14-3-3β at the actin cytoskeleton plays a role in maintaining the foot processes and the Par complex, and 14-3-3σ at the vimentin cytoskeleton is essential for maintaining the primary processes in podocytes, indicating that 14-3-3 proteins may serve as therapeutic targets for nephrotic syndromes.

## Materials and Methods

### Animals

Specific pathogen-free 7 weeks-old female Wistar rats (Charles River Japan, Atsugi, Japan) were used. All animal experiments conformed to the National Institutes of Health Guide for the Care and Use of Laboratory Animals. Procedures for the present study were approved by Animal Committee at Niigata University School of Medicine (Niigata, Japan; permit numbers SA01228), and all animals were treated according to the guidelines for animal experimentation of Niigata University.

### Cell culture of podocyte

Human cultured podocyte was donated by Dr. Moin A. Saleem (University of Bristol, Bristol, UK) (53). Mouse cultured podocyte was donated by Dr. P. Mundel (Albert Einstein College of Medicine, Bronx, NY) (54). Cultivation of conditionally immortalized mouse and human podocytes was conducted, as reported previously. A conditionally immortalized mouse and human podocyte cell line demonstrated nephrin and podocin expression. Podocytes were maintained in RPMI 1640 medium (Nissui Pharmaceutical, Tokyo, Japan) supplemented with 10% fetal bovine serum (FBS) (Life Technologies, Grand Island, Japan). For propagation of podocytes, cells were first cultivated at 33°C and maintained for 2 weeks at 37°C to induce differentiation. Human cultured podocytes were cultured with ADR 5 μg/ml for 24 hours and then harvested for immunofluorescence (IF), reverse transcription-polymerase chain reaction (RT-PCR), and western blot. Morphological analysis was performed by counting at least 100 cells in the full-frame images of each experiment, using image J software.

### RNA silencing analysis

The siRNA sequences targeting human 14-3-3β (GenBank accession number: NM_003404.4) and human 14-3-3σ (GenBank accession number: NM_006142.5) were synthesized by ThermoFisher Scientific Ink (Massachusetts, USA). The sense and antisense strands of siRNA were used. The sequence of the sense strand was 5’-GGGCAAAGAGUACCGUGAGAAGAUA-3’, corresponding to regions in exon 3 of YWHAB (14-3-3β). The sequence of the sense strand was 5’-GCGCAUCAUUGACUCAGCCCGGUCA-3’, corresponding to regions in exon 3 of SFN (14-3-3σ). BLAST searches of selected sequences revealed no significant homology to other human genes. Negative control siRNA of nonspecific nucleotide sequence was purchased from ThermoFisher Scientific Ink. Before treatment, human cultured podocytes and HEK293 cells were cultured to a density of 70-80% at 37°C, and then they were treated with the siRNA using HiPerFect Transfection Reagent (QIAGEN Inc), according to the manufacturer’s instructions. Cells were harvested for 72 hours after siRNA treatment for RT-PCR, IF, western blot, and IP analyses.

### Induction of Adriamycin (ADR)-induced nephropathy and Puromycin nucleotide (PAN)-induced nephropathy in rats

ADR-induced nephropathy was prepared by intravenous injection with 6 mg/kg BW. PAN-induced nephropathy was prepared by intravenous injection with 100 mg/kg BW, basically according to the method described previously (55, 56). The rats were sacrificed, and the kidneys were removed. The materials were used for IF, RT-PCR, and western blot. Small tissue samples were used for IF studies. To prepare a set of glomerular RNA for each time point, glomeruli were isolated from the remaining kidney tissue pooled from three rats each by a sieve method, and glomerular RNA was prepared with Trizol (Life Technologies, Inc., Gaithersburg, MD).

### Reverse transcription-polymerase chain reaction (RT-PCR) and real-Time RT-PCR

A semiquantitative RT-PCR analysis and real-time PCR analysis with the RNA of rat glomeruli and mouse and human cultured podocytes were performed basically according to the method described previously (57, 58). The sequences of the primers used for this study are provided in Supplemental Table S1. For semiquantitative RT-PCR, the band intensity was determined by image analysis using Bio Doc-It System and densitometry software, Lab Works 4.0 (UVP Inc., Upland, CA, USA). The results were corrected for the amount of mRNA in the sample by dividing by the intensity of the internal control glyceraldehyde-3-phosphate dehydrogenase (GAPDH). For real-time PCR, cDNA and specific primers were mixed with a TB Green Premix Ex Taq (Takara, Otsu, Japan). PCR reactions were run on a Smart Cycler System (Takara). Ct values of the gene targets were normalized to GAPDH. Fold change in the expression of target genes compared with the control was calculated using the 2-^ΔΔ^CT method with GAPDH. The sequences of the primers are shown in Expanded View Table 1.

### Immunofluorescence (IF) analysis

IF was performed basically according to the method previously reported (32, 33, 59–63). The following antibodies were used as the glomerular cell markers, mouse anti-rat endothelial cell antigen-1 (RECA-1) antibody, mouse monoclonal antibody to Thy1.1 (OX7) (60), mesangial cell marker, mouse monoclonal antibody to nephrin, slit diaphragm marker (mAb 5-1-6), mouse monoclonal antibody to podocalyxin, a podocyte apical marker, mouse monoclonal antibody to α3-integrin, and a podocyte basal marker, mouse monoclonal antibody to synaptopodin, a podocyte foot process marker. The antibody against FKBP12 was prepared in a rabbit immunized with a specific peptide (28). The suppliers of other antibodies used in this study are provided in Supplemental Table S2. To detect actin fiber in cultured podocytes, Rhodamine-Phalloidin was used according to the method of the previous report (32). To evaluate the 14-3-3β and 14-3-3σ staining in the rat glomeruli, the staining was graded as follows: continuous staining of >75% was score 4; 75-50% was score 3; 50-25% was score 2; and 25-0% was score 1. The intensity of staining was quantified by image analysis using Image J software (63, 64). A score and an intensity of staining were assigned to each glomerulus, and 30 glomeruli of each rat were analyzed using Image J software. A score and an intensity of staining were assigned to each cell, and at least 100 cells of each experiment were analyzed using Image J software. The data are shown as mean ± standard deviation (SD) (61).

### Western blot analysis

Western blot analysis was performed basically according to the method described previously (65, 66). In brief, rat glomeruli and human cultured podocytes were solubilized with sodium dodecyl sulfate (SDS)-polyacrylamide gel electrophoresis (PAGE) sample buffer (2% SDS, 10% glycerol, 6% mercaptoethanol in 62.5 mmol/L Tris-HCl [pH6.8] with protease inhibitors). The solubilized material was subjected to a polyvinylidene fluoride transfer membrane (Pall Corporation, Pensacola, FL). After exposure to the primary antibodies, alkaline phosphatase-conjugated secondary antibodies were used. The reaction was developed with an alkaline phosphatase chromogen kit (Biomedica, Foster City, CA). The band intensity was quantified by image analysis using Image J software.

An antibody list and primer information are described in Expanded View Table 1 and Table 2, respectively.

### Assay with HEK293 cell transfection and immunoprecipitation (IP) assay

Assays with HEK293 cell transfection were performed as previously described (59, 67). HEK293 cells were transfected with pFlagCMV2-14-3-3sigma (Addgene, Cambridge, MA, Plasmid #12453) (68), mCherry-vimentin7 (Addgene, Plasmid #55156) (69), pK-myc-Par3b (Addgene, Plasmid #19388) (70), a monomeric red fluorescent protein (RFP)-FKBP12 (Addgene, Plasmid #67514) (71) and synaptopodin-HA by the calcium phosphate method. The coding sequence of full-length synaptopodin was generated by PCR and subcloned into the pKH3 vector (Addgene). IP assays were performed basically according to the method described previously (59, 67). The cells were solubilized with 1% Triton X-100 or radioimmunoprecipitation assay (RIPA) buffer. The solubilized materials were analyzed by IP analysis.

### Statistical analyses

Statistical significance was evaluated using the unpaired *t*-test or Mann-Whitney *U*-test. Values were expressed as the mean ± standard deviation (SD). Differences at *P*<0.05 were considered significant. Data were analyzed using Graphpad Prism 5.0 software (Graphpad Software, San Diego, CA, USA).

## Acknowledgments

The authors wish to thank Ms. Mutsumi Kayaba for her excellent technical assistance. This work was supported by Grant-Aids for Scientific Research (22H03086 to HK and 20K08587 to YF) and Grant-Aid for Young Sciences (22K16235 to HY) from the Ministry of Education, Culture, Sports, Science and Technology of Japan.

## Expanded view figure legends

**Figure EV1 - Expression of 14-3-3 isoforms in the cultured podocytes.**

A mRNA expression of 14-3-3β, 14-3-3σ, and 14-3-3γ in undifferentiated and differentiated mouse cultured podocytes and mouse cerebrum. Although 14-3-3β, 14-3-3σ, and 14-3-3γ expression were detected in the mouse cerebrum, mRNA expression of 14-3-3σ was lower than that of 14-3-3β and 14-3-3σ. 14-3-3β and 14-3-3σ mRNA expression in the differentiated podocytes were higher than that in the undifferentiated cells. 14-3-3γ mRNA expression in the differentiated podocytes was lower than that in the undifferentiated cells. GAPDH was analyzed as the internal control.

B mRNA expression of 14-3-3ε, 14-3-3ζ, 14-3-3η, and 14-3-3θ in undifferentiated and differentiated mouse cultured podocytes and mouse cerebrum. 14-3-3ε, 14-3-3ζ, 14-3-3η, and 14-3-3θ mRNA expression were detected in the mouse cerebrum. Alteration in these mRNA expressions was not detected in the differentiated and the undifferentiated cells.

C mRNA expression of 14-3-3β and 14-3-3σ in undifferentiated and differentiated human cultured podocytes. 14-3-3β and 14-3-3σ mRNA expressions in the differentiated cells were higher than that in the undifferentiated cells.

D Protein expression of 14-3-3β and 14-3-3σ in undifferentiated and differentiated human cultured podocytes. Protein expression of 14-3-3β and 14-3-3σ in the differentiated cells was significantly higher than that in the undifferentiated cells. Actin was analyzed as the internal control. (*n*=3; *p<0.05, ***p<0.001, *t*-test)

**Figure EV2 - Localization of vimentin and synaptopodin in kidneys.**

Vimentin staining was detected in the glomeruli and the tubular interstitial cells. Synaptopodin staining was restricted in the glomeruli. In glomeruli, vimentin staining was clearly apart from synaptopodin. Scale bar, 20 μm.

**Figure EV3 - Proportion of the cells with stress fibers and vimentin fibers in 14-3-3β knockdown (KD) cultured podocytes and 14-3-3σ KD cultured podocytes.**

A Proportion of the cells with only stress fibers and the cells without stress fibers and vimentin fibers in the cultured podocytes treated with 14-3-3β siRNA. Proportion of the cells with only stress fibers in the cytoplasm was not altered by the treatment of 14-3-3β siRNA (left panel). Proportion of the cells without stress fibers and vimentin fibers was not altered by the 14-3-3β siRNA (right panel). (*n*=3-4; N.S., non-significant, *t*-test)

B Proportion of the cells with only vimentin fibers and the cells without stress fibers and vimentin fibers in the podocytes treated with 14-3-3σ siRNA. Proportion of the cells with only vimentin fibers was decreased by the treatment of 14-3-3σ siRNA (left panel). Proportion of the cells without stress fibers and vimentin fibers was not altered by the 14-3-3σ siRNA (right panel). GAPDH was analyzed as the internal control. (*n*=3-4; ***p<0.001, N.S., non-significant, *t*-test)

**Figure EV4 - Localization of 14-3-3σ in glomeruli of ADR-induced nephropathy and mRNA expression of 14-3-3σ in the cultured podocyte treated with ADR.**

A localization of 14-3-3σ and podocalyxin on day 28 of ADR nephropathy. 14-3-3σ stating was clearly apart from podocalyxin in ADR nephropathy. The remaining 14-3-3σ staining was co-localized with Thy.1.1. Scale bar, 20 μm.

B 14-3-3σ mRNA expression was increased in the cultured podocytes treated with ADR for both 14h and 24h. GAPDH was analyzed as the internal control. (*n*=3-4; **p<0.01, ***p<0.001, *t*-test)

